# Persistent SARS-CoV-2 Spike is Associated with Localized Immune Dysregulation in Long COVID Gut Biopsies

**DOI:** 10.64898/2026.03.09.707564

**Authors:** Salim Abraham Soria, Patrick Peterson, Michael Tankelevich, Saurabh Mehandru, David Putrino, Marcelo Freire

## Abstract

SARS-CoV-2 persistence is a proposed driver of Long COVID (LC), but the *in-situ* relationship between residual viral antigen and immune dysregulation remains poorly defined. To address this critical gap, we employed a high-resolution, multi-modal approach—combining RNAscope, GeoMx Digital Spatial Profiling (DSP), spatial transcriptomics, and multiplex immunofluorescence—on 25 terminal ileum and left colon biopsies from a clinical cohort of 8 LC participants and 5 healthy controls. We confirmed the persistence of SARS-CoV-2 Spike transcript and protein in the gut tissue of all LC cases and controls tested. Yet, comparison of Spike-positive (Spike+) regions in LC versus healthy control colon tissues revealed a differential, symptomatic state-associated signature, with 57 differentially expressed genes (DEGs) (26 upregulated, 31 downregulated), revealing genes that disrupt the immune response in LC subjects. LC colon Spike+ regions demonstrated increased expression of AQP8 and other absorptive-related genes (SLC26A3, SLC26A2, and CLCA4) which are involved with Chron’s disease along with transcripts involved in tumorigenesis (GUCA2A, S100P, TSPAN1). Simultaneous downregulation of key homeostatic chemokines (*CXCL13, CCL19, CCL21*), and other transcripts reported to exhibit low expression in colorectal cancers (TMEM88B, NIBAN3, DMBT1), suggesting a paradox of epithelial tissue stress yet dysfunctional immune trafficking. Further analysis comparing Spike+ versus Spike-regions *within* LC colon tissue demonstrated an active, localized, antigen-driven immune microenvironment, identifying 122 DEGs (82 upregulated, 40 downregulated), including tumorigenesis genes. Cellular deconvolution of Spike+ regions revealed a statistically significant focal enrichment of myeloid-derived cells (macrophages, non-classical/intermediate monocytes), plasma cells, and regulatory T cells, coupled with significant enrichment in T-cell-related pathways, including “Antigen processing and presentation,” and “Th1/Th2/Th17 cell differentiation.” The ileum displayed a similar, though less pronounced, signature, demonstrating these statistically significant findings are specific to the colon of LC subjects. In contrast, corresponding Spike+ vs. Spike-analysis in healthy control colon tissues showed a more modest transcriptional response with 38 DEGs. Our data provide robust evidence that persistent SARS-CoV-2 Spike protein detection in the gut is not immunologically inert. Instead, it is actively associated with distinct, immune cell composition shifts and a dysfunctional pro-inflammatory transcriptional profile, supporting the hypothesis that retained viral antigen drives chronic immune dysregulation in tissue of Long COVID subjects.

## Introduction

Long COVID (LC), an Infection-Associated Chronic Condition (IACC) triggered by acute SARS-CoV-2 infection, was reported by 6.9% of U.S. adults in 2022 as having ever experienced the condition.^1^ The WHO defines Long COVID as a chronic disease arising from SARS-CoV-2 infection, with symptoms persisting beyond 3 months post-infection.^2,3^ LC presents with marked symptom heterogeneity, spanning multiple organ systems and clinical phenotypes.^4^ However, epidemiology studies have characterized the most common symptoms as: post-exertional malaise, fatigue, dizziness, cognitive impairment, and gastrointestinal symptoms.^5–7^ LC rarely resolves spontaneously; most patients endure debilitating symptoms that substantially diminish quality of life relative to healthy individuals and those with other chronic conditions.^8–11^ A 2022 U.S. study reported that approximately 7.5% of Americans experienced Long COVID, and affected individuals averaged about 8.4 days of missed work because of illness or injury compared to 5.2 and 3.1 days among individuals with prior COVID-19 infection without persistent symptoms and those who never had COVID-19, respectively.^12^ Although there are efforts to treat and manage symptoms, no specific drug or treatment has been identified that may target any of the potential root causes of Long COVID.^13^ Outside the clinic, the total U.S. economic burden of Long COVID is estimated to reach $3.7 trillion in lost quality of life, income, and additional healthcare-related costs.^14,15^ The complexity of LC and its devastating burden demand urgent mechanistic research into its pathogenesis.

Since the pandemic began, initial studies have looked at how the presence of SARS-CoV-2 affects the local immune and parenchymal cellular environment of patients with an active COVID-19 infection. A 2021 autopsy study detected SARS-CoV-2 RNA within the lungs of patients who had succumbed to SARS-CoV-2 and characterized the immune and alveolar cellular environment.^16^ This report did not detect the presence of viral RNA in other tissue types (heart, liver, and kidney) collected that are high expressors of the SARS-CoV-2 entry factors (ACE2 and TMPRSS2). However, a second study detected the presence of the SARS-CoV-2 Nucleocapsid protein in the duodenum and/or ileum of patients with acute COVID-19.^17^ More specifically, the nucleocapsid protein was detected broadly in intestinal epithelial cells (high expressors of ACE2).^18^ Other studies also report the detection of SARS-CoV-2 RNA in stool during acute infection.^7,19^ Research on acute COVID-19 infections have shown that viral RNA or protein can be found in other organs beyond the primary site of infection in the respiratory tract. Mounting evidence now supports viral RNA or protein persistence as a driver of LC symptoms.^20^ Multiple studies report SARS-CoV-2 RNA or protein in blood, stool, and tissue obtained from Long COVID patients.^20–26^ Among biofluids and tissues explored, the gastrointestinal tract has emerged as a site of particular interest in LC. Its vast mucosal surface, intimate association with the gut microbiome, and dense immune surveillance make it an ideal niche for persistent viral antigen.^27^ In 2024, a study was published showing the detection of viral RNA persistence in the colon for up to 2 years since SARS-CoV-2 infection in patients with reported Long COVID symptoms.^24^ Together with acute COVID-19 reports, these findings implicate the lower gastrointestinal tract as a key site driving LC pathogenesis and chronicity.

While the evidence of SARS-CoV-2 RNA or protein persistence is strong^20,24,28^, a critical gap remains in understanding the localized, *in-situ* biological impact in Long COVID. How viral RNA and proteins reshape the local immune microenvironment to drive sustained pathology remains unknown. To close this gap, we employed a high-resolution, multi-modal approach—RNAscope, spatial transcriptomics (GeoMx DSP), and multiplex immunofluorescence—on terminal ileum and left colon biopsies from LC participants and healthy controls to define the spatially-resolved immune response to persistent viral antigen.

## Materials and Methods

### Ethics Statement and Inclusion Criteria

All study procedures were approved by Mount Sinai’s Program for Protection of Human Subjects (STUDY# 16-00512). All participants signed an informed consent document before enrollment into the study. Study participants were either healthy controls who were receiving a routine colonoscopy or people who were diagnosed with Long COVID by a clinical provider according to the Centers for Disease Control and Prevention and National Academies of Science, Engineering, and Medicine guidelines.^2^ Enrollment of participants occurred over the years of 2021 to 2025. Demographics were collected and included infection history, vaccination history, comorbidities, and medications (**Supplementary Table 1, 2, 3**). People with Long COVID were stratified into those who reported persistent gastrointestinal symptoms and those who did not.

### Study Participants

Thirteen participants were recruited in this study, consisting of 8 (6 female) Long COVID and 5 (1 female) controls to produce 25 total samples (13 colon and 12 ileum) (**Supplementary Table 1)**. One LC participant was collected twice at different time points, and a different LC participant had only their left colon sampled. From the Long COVID cohort, 5 participants report GI comorbidities/ symptoms while 3 participants do not (**Supplementary Table 2**). Samples from LC participants were collected between 7 months and 2.5 years following their last reported SARS-CoV-2 infection. All participants, except for 1 LC and 1 control, received 1 or more vaccinations against COVID-19 (**Supplementary Table 3**).

### Sample Collection

All participants underwent an upper GI endoscopy/colonoscopy as previously reported.^17^ In brief, intestinal biopsy tissues derived from the terminal ileum and left colon were collected into 10% neutral-buffered formalin (NBF) and fixed for 24-48 hours. Afterwards, the NBF solution was replaced by 70% isopropanol and shipped to JCVI (La Jolla, CA). Samples were processed into formalin-fixed paraffin-embedded (FFPE) tissue blocks, then sliced for H&E staining and serial sections to use in the GeoMx DSP RNA FFPE protocol (MAN-10150-01). Serial sections of tissue were cut at 5 µm thickness and mounted on slides following constraints for GeoMx DSP use.

### H&E Staining

Hematoxylin & Eosin (H&E) staining was performed through the histology core at La Jolla Institute for Immunology using a Leica ST5020 Automated multistainer. Briefly, slides were baked twice for 5 minutes, followed by 3 washes of Pro-Par at 5 minutes each, 2 washes of 100% isopropanol, and 1 wash of 90% isopropanol. SelectTech 560 Hematoxylin (Leica Biosystems Ca. #3801571) and Eosin Phloxine 515 SelecTech (Leica Biosystems Ca. #3801606) were used to stain tissues with SelecTech Define (Leica Biosystems Ca. #3803591) and SelecTech Blue Buffer 8 (Leica Biosystems Ca. #3802916) buffers to improve staining quality. H&E-stained tissues were imaged using the Zeiss Axioscan Z1 slide scanner at 40x objective, then .czi image files were opened on QuPath (v6.0) and visualized.

### RNAscope Staining

To detect the presence of viral RNA, RNAscope staining targeting key transcripts was performed on collected tissue using the RNAscope Multiplex Fluorescent Reagent Kit v2 protocol (UM 323100) followed by immunofluorescent staining then imaged with the GeoMx DSP. The key transcriptional targets are the antisense spike gene (*nCoV2019-S*) and the positive sense open reading frame 1ab (*nCoV2019-orf1ab*-sense) gene. RNAscope probes used to detect these transcripts were V-nCoV2019-S (Advanced Cell Diagnostics Ca. 848561; target region 21,631 - 23,303) and V-nCoV2019-orf1ab-sense (Advanced Cell Diagnostics Ca. 859151-C2; target region 1,583 - 4,388). Additionally, positive and negative technical slides were run using probes targeting human housekeeping genes (*POLR2A* and *PPIB*) and *dapB*, respectively.

Slides were baked overnight at 60°C in a dry oven, followed by deparaffination and rehydration using two washes of xylene at 5 minutes each and two washes of 100% ethanol at 2 minutes each. Slides were then dried at 60°C in an oven until all excess ethanol had evaporated. The slides were then incubated with hydrogen peroxide for 10 minutes at room temperature (RT), washed twice, then treated with target retrieval for 15 minutes at >99°C using a Hamilton Beach digital steamer (37530Z). A hydrophobic pen (Vector Labs, Ca. 310018) was used to create a hydrophobic barrier surrounding the tissue. Once the barrier was dried, RNAscope Protease Plus was added to the slide and incubated for 30 minutes at 40°C using the RapidFISH Slide Hybridizer (Boekel Scientific), followed by two washes. A probe mix solution was made by mixing V-nCoV2019-S-C1 and 50x V-nCoV2019-orf1ab-sense to produce a final solution where both probes are at a 1x concentration. Slides were then incubated with this probe mix solution, positive technical control probes, or negative technical control probes for 2 hours at 40°C, washed twice, then left overnight in 5x SCC at RT. To begin signal amplification, tissues were hybridized with RNAscope Multiplex FL v2 Amp1, Amp2, and Amp3, then signal was developed using HRP-C1 coupled to TSA Vivid Fluorophore 570 (conc. 1:1500; ACD 323272) and HRP-C2 coupled to TSA Vivid Fluorophore 650 (conc. 1:1200; ACD 323273) as described in RNAscope Multiplex Fluorescent Reagent Kit v2 protocol (UM 323100). Following the last HRP blocking step, slides were incubated with a morphology mix containing SYTO-13 [ThermoFischer, Ca. S7575] and anti-Pan-CK [Novus; Ca. NBP2-76425AF594] for 1 hour at RT. Slides were washed twice for 5 minutes using 2x SCC, then imaged using the GeoMx DSP set to “Scan Only”.

### Spatial Transcriptomics

To perform Spatial Transcriptomics, the GeoMx DSP Manual Slide Preparation (MAN-10150-01) was followed. Briefly, slides were baked in an oven at 60°C overnight. The slides were then transferred into Simportx^TM^ Scientific EasyDip^TM^ slide staining jars (FisherScientific 22-038-489) and deparaffinized by 3 washes of xylene at 5 minutes each, followed by rehydration with two washes of 100% EtOH, then 95% EtOH at 5 minutes each. Slides were transferred to 1x PBS for 1 minute. Slides were then transferred to a jar containing 1x PBS at >99°C for 10 seconds and immediately transferred to 1x IHC antigen retrieval solution (Invitrogen Ca. 00-4956-58) at >99°C for 15 minutes using a Hamilton Beach digital steamer (37530Z). Slides were removed and placed in 1x PBS to cool down for 5 minutes, then transferred to a jar containing 1x PBS with 1 µg/mL Proteinase K (Invitrogen AM2548) at 37°C for 15 minutes. Afterwards, another 1x PBS wash was performed for 5 minutes, then immediately transferred to 10% NBF for 5 minutes, followed by two washes of NBF Stop buffer each at 5 minutes and 1x PBS at 5 minutes.

*In-situ* hybridization was performed by adding a probe mix solution containing 200 µL of Buffer R, 25 µL of Human Whole-Transcriptome (Hu WTA) probe mix, and 25 µL of DEPC-treated H_2_O per slide. All materials following incubation with the probe mix solution were cleaned using RNase AWAY (ThermoFisher 7003PK). Slides were removed from 1x PBS and transferred to a slide tray, where ∼200 µL of the probe mix solution was added to each slide, and a coverslip (Grace Bio-Labs HybriSlip) was set on top. The slide tray was then placed inside the RapidFISH Slide Hybridizer (Boekel Scientific) for an overnight incubation at 37°C. After the incubation, slides were dipped in 2X SSC until the coverslips slipped off, then washed twice in a mixture of 50:50 4x SSC and 100% formamide (Ambion Ca. AM9343) at 37°C for 25 minutes each, followed by two washes of 2x SSC at 2 minutes each. Slides were then transferred to a humidity chamber and blocked with ∼200 µL of Buffer W (Bruker/Nanostring) for 30 minutes, protected from light. While waiting, a morphology marker solution was made mixing 1:1000 dilution of SYTO-13 (Invitrogen Ca. S7575), anti-SARS-CoV-2 Spike (Novus; Ca. NBP2-90980AF532), anti-PanCK (Novus; Ca. NBP2-76425AF594), and anti-CD19 (Novus; Ca. NBP2-25196AF647) at 2µg per 200µL. After blocking and removing excess Buffer W, the morphology marker solution was added until all tissues were covered (∼200µL), then incubated at RT for one hour. Lastly, the slides were washed twice in 2x SSC and then loaded onto the GeoMx for imaging and ROI collection.

Based on previous “Scan Only” runs, an ROI collection strategy was developed based on the presence of SARS-CoV-2 Spike protein or CD19 signal. In the presence of a SARS-CoV-2 Spike protein signal, a circular ROI was created and labeled as Spike+ ROIs. Regions without any Spike protein signal were labeled as Spike-. In regions with CD19^+^ cell signal, circular ROIs were created where CD19+ cells were found without the presence of Spike protein signal. ROIs labeled CD19-were selected based on the negative signal of both Spike protein and CD19. For each of these ROI classes, three ROIs were selected per patient’s colon or ileum sample to a total of 24 ROIs per participant or 252 total ROIs for the study (132 for colon and 120 for ileum). After oligo barcode collection of each ROI, the collected oligos were processed for NGS sequencing using custom NGS primers developed for GeoMx DSP use (MAN-10153-01). Pooled libraries were sequenced using the NovaSeq 6000 platform and a SP flow cell. FASTQ files and the GeoMx DSP configuration file (.ini) were transferred to Illumina’s Basespace platform and processed through the GeoMx NGS Pipeline to produce a digital count conversion (.dcc) file. Lastly, the DCC file was transferred back to the GeoMx DSP for a final demultiplexing step to produce the working dataset.

### Multiplex Immunofluorescence

Serial sections of the tissue samples (colon and ileum) used in the spatial transcriptomics run were processed by Akoya Biosciences as part of a proof-of-concept program utilizing the PhenoCycler-Fusion system. This ultrahigh-plex imaging platform was employed with a 27-plex custom immune panel targeting key immune and cellular markers relevant to the gut microenvironment.

The tissue sections were stained with oligonucleotide-barcoded antibodies in a single step, followed by iterative imaging cycles. In each cycle, fluorescent reporters complementary to a subset of the antibody barcodes were flowed onto the tissue, imaged, and then chemically removed. This sequential process allowed for the precise spatial mapping of over two dozen markers on the same tissue section without spectral overlap. The resulting single-cell resolution protein data were generated for subsequent co-registration and integrated analysis with the corresponding spatial transcriptomics data, enabling deep cellular phenotyping and the contextualization of gene expression within the intact gut tissue architecture.

### Analysis & Bioinformatics

All images produced by the GeoMx DSP were exported as OME.TIFF files were further processed in QuPath (v6.0). Within QuPath, all images were processed through a cell detection and feature extraction pipeline. Briefly, a pixel classifier was trained to create annotations that contain only the area the tissue covers. Cell detection was performed, and intensity features were calculated for all channels per cell. Single-cell fluorescent intensity data were then exported from QuPath into R (R version 4.5.1). *Tidyverse* was used for data processing, then plotted using *ggplot2* coupled with *ggpubr* to produce box-plots and volcano-plots with statistics overlaid.^29^

The resulting spatial transcriptomic dataset was exported from the GeoMx DSP and imported into R. The initial dataset was converted into a *standR* object processed through quality control checks, filters, and upper-quantile normalized.^30^ Differential expression analysis was performed using the *limma-voom* pipeline. The model matrix was designed without an intercept, the condition of the participant, and the marker for which the ROI is positive, while also considering the percentage of cells positive for Spike protein within the ROI (*∼ 0 + Status: Marker + Total_Spike_pct*). Additionally, correlation and blocking terms were added to account for multiple ROIs belonging to the same patient. Log_2_FC and p-values (uncorrected and Benjamini-Hochberg corrected) were saved by contrast to create visualizations and further downstream analysis.

Spatial deconvolution of ROI data was performed using *SpatialDecon* coupled with the safeTME, a cell identity dataset heavily annotated with immune cell types, following an established bioinformatic pipeline (*propeller framework*).^31^ Violin plots with annotated FDR statistics were made using ggplot. Gene-set-enrichment analysis (GSEA) was performed using *clusterProfiler* with *gseKEGG* and *gseGO* annotated with *org.Hs.eg.db*.^32^ The dataset of all genes after differential expression was organized in descending log_2_FC value and annotated with EntrezIDs. Dotplot and emapplot visualizations were made using *clusterProfile*. Results produced by *gseGO* were simplified to remove redundant or highly overlapping gene sets using *simplify()* (cutoff = 0.7).

Phenocycler-fusion multiplex immunofluorescent data were analyzed using QuPath (v6.0). An object classifier was made to detect positive cells from the background and annotate cells based on what markers they express. Single-cell fluorescent data was processed using an established bioinformatic pipeline in R.^33^ Briefly, immunofluorescent data were z-score normalized and batch corrected using *removeBatchEffect* (*limma*). Principal component analysis (PCA) was performed on immunofluorescent data by tissue type followed by a kNN graph of the PCA (k = 20), then clustered based on neighbors to annotate cells with detected Leiden clusters and ultimately plot each cell on a UMAP. *DittoSeq* was used to produce a dotplot that plots “relative expression” and “% Expressing”.^34^ “Relative expression” represents the median expression of all cells with a non-zero fluorescent intensity for one cluster and scaled against all clusters for one marker. “% Expressing” represents the number of cells in a cluster that were marked as positive by the QuPath classifier, divided by the total number of cells for that cluster. Leiden ID annotations were then imported into QuPath using a custom script that colors detected objects by the same color used on the UMAP colored by Leiden ID.

## Results

### Long COVID versus a Healthy baseline

H&E-stained CZI images were analyzed via QuPath (v6.0) using a pixel classifier to determine tissue area, along with a cell detection and classification pipeline to obtain “Epithelium” and “Stroma” cell classes (**Figure 1B**). **Figure 1C** plots the results of these measurements by grouping the number of cells per mm^2^ of tissue for each tissue compartment to compare Long COVID against Healthy samples. No statistical significance was observed when comparing the number of detected cells per mm^2^ of tissue using a Mann-Whitney U Test (Colon/ Epithelium p = 0.921, Colon/ Stroma p = 0.776, Ileum/ Epithelium p = 0.667, and Ileum/ Stroma p = 0.833).

**Figure 1.**
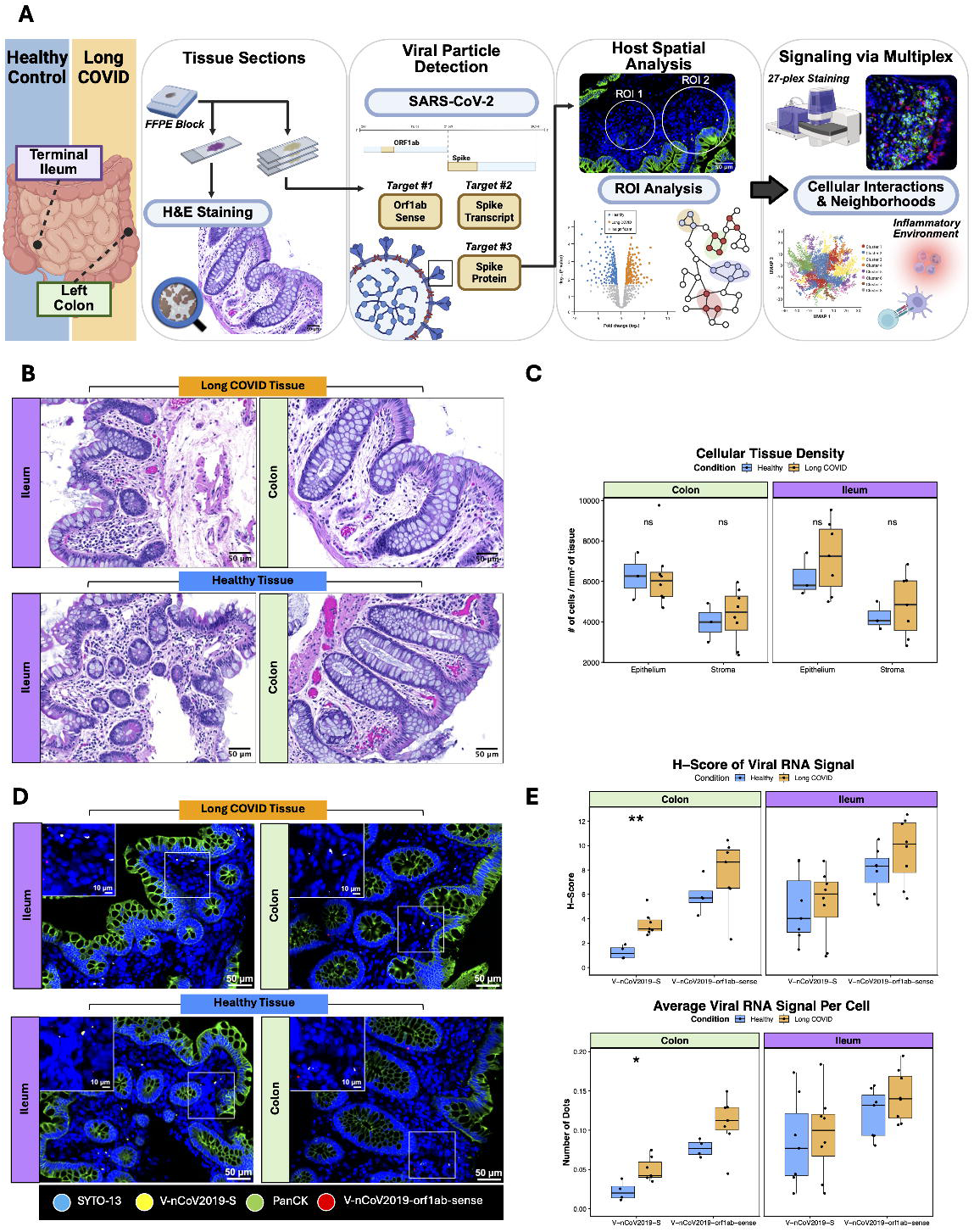
Summary of multi-modal approach and initial tissue findings. (A) Summary of methods and pipelines (13 participants; Long COVID = 8; Healthy = 5). (B) Representative images of H&E stains for both the terminal ileum and the left colon. For each tissue type, either 2 or 3 replicate biological samples were collected (C) A QuPath pixel classifier was trained to detect tissue and produce an estimate for area (Long COVID = 8; Healthy = 3). A cell detection and classification pipeline was made to differentiate between epithelium and stroma cells. An initial quantification of cell density was measured between Long COVID and Healthy samples separated by tissue and type of cell. A Mann-Whitney U test was performed, and no statistically significant differences were detected (*p* ≤ 0.05 (*), *p* ≤ 0.01 (**), *p* ≤ 0.001 (***)) (D) Representative immunofluorescent images stained with RNAscope probes (*V-nCoV2019-S* and *V-nCoV2019-orf1ab-sense*) stratified by tissue type (Ileum or Colon) and condition (Long COVID or Healthy). White arrows point at *V-nCoV2019-S* signal. (E) QuPath was used to detect punctuate probe signals, and results were summarized as either an *H-Score* or *Signal per Cell* (Long COVID = 4; Healthy = 2). Data were stratified by tissue type and RNAscope probe. Comparison of *V-nCoV2019-S* signal in Colon saw a statistically significant difference in both *H-Score* and *Average Signal per Cell* based on a Mann-Whitney U test (*p* ≤ 0.05 (*), *p* ≤ 0.01 (**), *p* ≤ 0.001 (***), p ≤ 0.0001 (****)).

To detect the presence and location of where viral RNA persists, RNAscope probes V-nCoV2019-S and V-nCoV2019-orf1ab-sense alongside anti-PanCK were used in a dual RNAscope/immunofluorescent protocol in ileum and colon (**Figure 1D**). **Figure 1E** depicts two different measurements to determine differences in the level of signal between tissues. When comparing the H-Score of tissues, a statistical significance (p = 0.006, Holm correction p = 0.012) was observed in the signal for the probe V-nCoV2019-S where the Healthy and Long COVID group have a median of 1.160 and 3.158, respectively. Unlike its colon counterpart, ileum tissues hybridized with V-nCoV2019-S had no statistically significant H-score difference between groups (Long COVID Median = 6.040, Healthy Median = 4.032; p = 0.867, Holm correction p = 0.87). No significant differences were observed in V-nCoV2019-orf1ab-sense signal within colon (Long COVID Median = 8.645, Healthy Median = 5.709; p = 0.164, Holm correction p = 0.16) or ileum tissue (Long COVID Median = 10.118, Healthy Median = 8.331; p = 0.232, Holm correction p = 0.46). Similarly, the average number of punctuate dots per cell was statistically significant for V-nCoV2019-S in colon (Long COVID Median = 0.042, Healthy Median = 0.020; p = 0.012, Holm correction p = 0.024), while ileum was not statistically significant (Long COVID Median = 0.100, Healthy Median = 0.077; p = 0.694, Holm correction p = 0.69). On the other hand, the orf1ab probe did not show any statistical significance in colon (Long COVID Median = 0.112, Healthy Median = 0.077; p = 0.073, Holm correction p = 0.073) and in ileum (Long COVID Median = 0.132, Healthy Median = 0.140; p = 0.189, Holm correction p = 0.38).

Next, we stained the tissues with an antibody panel targeting SARS-CoV-2 Spike protein, PanCK, and CD19 (**Figure 2A**). Tissue images were then analyzed via QuPath, where cells were detected and classified as positive or negative for each of the three antibody markers. **Figure 2B** shows the results of detected and classified Spike protein positive cells between the two tissue types and compartments for all Long COVID samples and 3 out of the 5 control samples. In colon tissue, no statistically significant difference in the percentage of cells showing positive signal was observed between Long COVID and controls in either the epithelium (Long COVID Median = 0.160, Healthy Median = 0.241; p = 0.921, Holm correction p = 0.99) or the stroma (Long COVID Median = 0.676, Healthy Median = 0.942; p = 0.497, Holm correction p = 0.99). Results for the ileum samples also show no statistically significant difference between groups in epithelium (Long COVID Median = 0.884, Healthy Median = 1.482; p = 0.267, Holm correction p = 0.53) and stroma (Long COVID Median = 2.184, Healthy Median = 1.588; p = 0.383, Holm correction p = 0.53).

**Figure 2.**
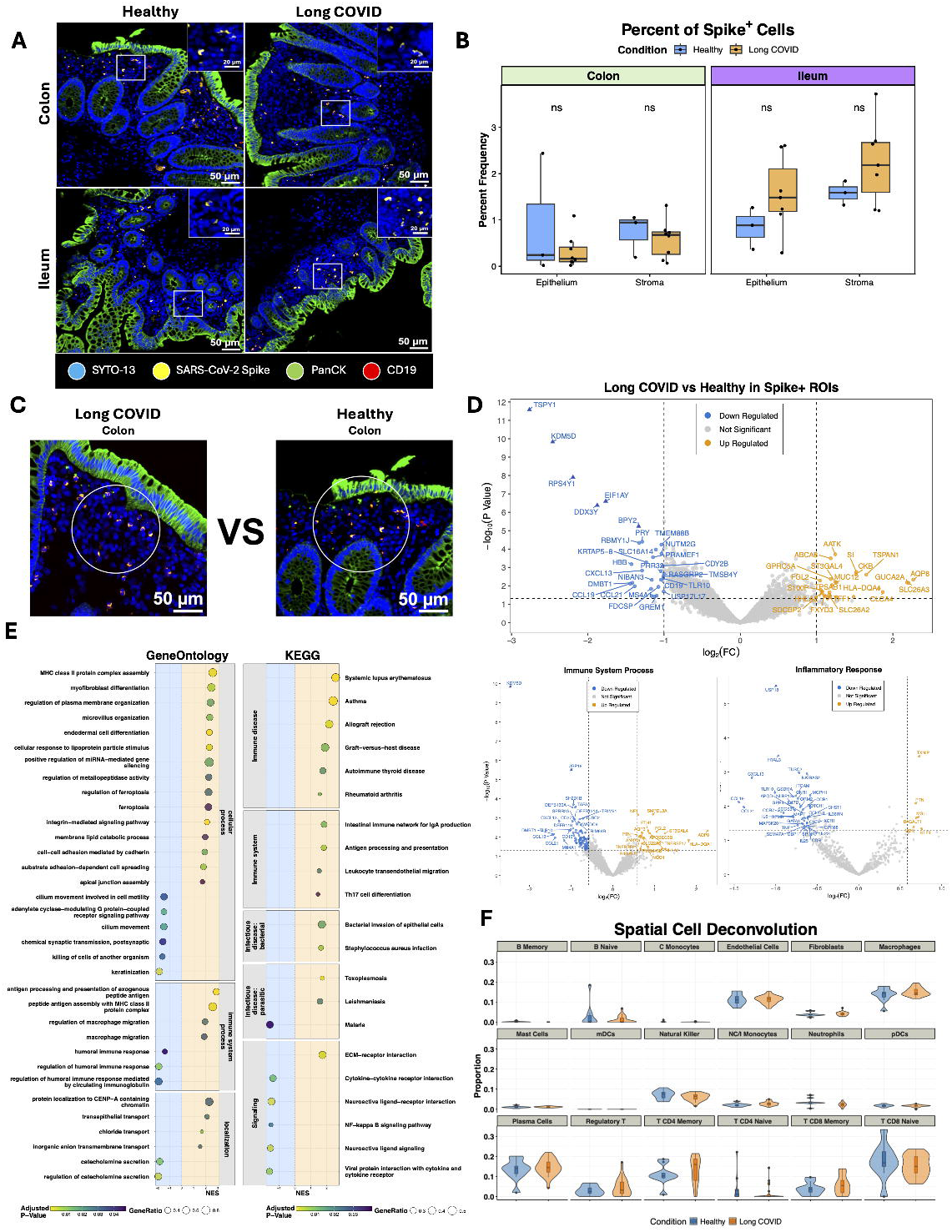
Long COVID and Healthy Immunological Differences by Spatial Transcriptomics. (A) Representative staining images of the left colon and terminal ileum between healthy and Long COVID participants used for ROI collection. Tissue sections were stained with SYTO-13 (Blue), anti-SARS-CoV-2 Spike (CR3022) (Yellow), anti-PanCK (Green), and anti*-CD19 (CB19)* (red) and visualized using the GeoMx DSP (Long COVID = 8; Healthy = 3). (B) SARS-COV-2 Spike protein positive cell detection based on a QuPath pipeline comparing condition and tissue type between tissue compartments. Cell detection was performed using an adapted StarDist script followed by cell classification based on a supervised ML model. Kruskal-Wallis with multiple comparisons was performed (*p* ≤ 0.05 (*), *p* ≤ 0.01 (**), *p* ≤ 0.001 (***)).90. (C) ROI selection and differential expression strategies comparing detected Spike protein positive ROI’s in colon tissue between Long COVID and healthy participants. (D) Differential analysis was performed using the limma-voom pipeline, correcting for replicate samples and variation in the frequency of detected SARS-CoV-2 Spike by ROI. 18,582 transcript targets were plotted. The main volcano plot highlights 26 and 31 genes that are upregulated and downregulated, respectively, at an unadjusted *p*-value threshold of 0.05 and an absolute log_2_FC > 1. Points with a triangular shape passed the adjusted p-value threshold based on a Benjamini-Hochberg correction. Smaller volcano plots show differentially expressed genes belonging to *GeneOntology* (GO) panels (GO:0002376 & GO:0006954) at an unadjusted *p*-value threshold of 0.05 and a Fold-Change value of 1.5. (E) Gene-Set Enrichment Analysis was performed on a ranked list of log_2_FC values with *GeneOntology* and *KEGG (*54 and 38 statistically significant pathways, respectively). Representative categories were selected, and pathways/ gene sets were plotted using the normalized enrichment score (NES) alongside the ratio of genes in the leading edge to total genes in the gene set (GeneRatio) and the adjusted p-value. (F) Spatial deconvolution was performed using a safeTME reference expression profile. Statistical significance was assessed using the propeller framework, a moderated two-tailed Mann-Whitney U test with BH FDR correction (*FDR* ≤ 0.05 (*), *FDR* ≤ 0.01 (**), *FDR* ≤ 0.001 (***)).

Based on the consistent detection of SARS-CoV-2 Spike protein positive cells using an antibody in both tissue types and in Long COVID and control participants, a GeoMx DSP project was designed to probe these cells and their surrounding environment for gene expression changes using regions-of-interest (ROIs) and the human whole transcriptome atlas (GeoMx Hu WTA) of 18,676 protein-coding genes. After initial processing and filtering, the limma-voom pipeline was used to perform differential analysis between Long COVID and control samples in Spike protein positive ROIs (labeled as Spike+) (**Figure 2C**). The volcano plot comparing Long COVID (n = 24 ROIs; 8 samples) and Healthy (n = 9 ROIs; 3 samples) Spike+ ROIs in colon tissue (18,582 transcripts) showed 31 downregulated transcripts and 26 upregulated transcripts for raw p-values < 0.05 and a log_2_FC threshold of |1| (FC = 2) (i.e., a 2-fold increase or decrease) (**Figure 2D & Supplementary Figure 2A**). When performing an adjustment for false discovery rate (Benjamini-Hochberg; BH) only 6 downregulated genes passed the p-adjusted threshold of <0.05 (p-adjusted values; *BPY2* p = 0.015, *DDX3Y* p = 0.0016, *EIF1AY* p = 0.0012, *RPS4Y1* p = 7.7x10^-5^, *KDM5D* p = 1.4x10^-6^, and *TSPY1* p = 4.9x10^-8^). Additionally, two separate volcano plots were made with less stringent FC thresholds (FC = 1.5) while selecting only genes that are found under the GO categories for immune system process (GO: 0002376) and inflammatory response (GO:0006954). In the immune system process volcano plot (total transcripts = 2555), 23 transcripts were upregulated, and 99 were downregulated based on raw p-values. When applying the BH correction, only 2 downregulated transcripts remained statistically significant (*USP18* p-adjusted = 9.6x10^-3^, and *KDM5D* p-adjusted = 1.4x10^-6^). The inflammatory response volcano plot had 6 and 43 transcripts upregulated and downregulated, respectively. A BH correction showed only 1 transcript statistically significant (*USP18* p-adjusted = 9.6x10^-3^).

Transcript log_2_FC values were ranked in decreasing order and fed through the *clusterProfile* pipeline to perform gene-set-enrichment-analysis (GSEA). Results were annotated using KEGG and GO biological knowledgebases. Genesets, of representative categories, were plotted with normalized enrichment score (NES) along with their BH-adjusted p-value and GeneRatio (defined as the ratio of leading-edge genes to total geneset size) (**Figure 2E**). KEGG GSEA thresholded to p-adjusted < 0.05 (BH corrected) identified 38 total statistically significant pathways, of which 29 were enriched (NES > 0) and 9 were downregulated (NES < 0). The most represented categories were Immune disease (6 pathways; all enriched), Signaling molecules and interaction (5 pathways; 1 enriched and 4 downregulated), and Immune system (4 pathways; all enriched) (**Supplementary Figure 2B & 2C**). Notable KEGG pathways include Intestinal immune network for IgA production, Antigen processing and presentation, and Bacterial invasion of epithelial cells, while downregulated pathways of interest are Viral protein interaction with cytokine and cytokine receptor, Neuroactive ligand signaling, and NF kappa B signaling pathway. GO GSEA indicated 36 enriched and 18 downregulated gene sets for a total of 54 statistically significant (p-adjusted < 0.05) gene sets. The top 3 categories from the first hierarchical level were cellular process (21 gene sets; 14 enriched and 7 downregulated), immune system process (7 genesets; 4 enriched and 3 downregulated), and multicellular organismal response (7 gene sets; 4 enriched and 3 downregulated) (**Figure 2E**). Spatial cell deconvolution was performed to determine the immune environment of ROIs that are SARS-CoV-2 Spike protein positive. Results between ROIs belonging to both Long COVID and Healthy were compared (**Figure 2F**). No statistically significant differences were observed in the proportion of genes attributable to any immune cell type after FDR p-value correction.

Differential analysis of the terminal ileum of Long COVID (n = 21 ROIs; 7 samples) and Healthy (n = 9 ROIs; 3 samples) Spike+ ROIs (18,581 transcripts) showed 10 downregulated transcripts and 8 upregulated transcripts for raw p-values < 0.05 and a log_2_FC threshold of |1| (FC = 2) (i.e., a 2-fold increase or decrease) (**Supplementary Figure 1A & 2D)**. However, only 3 downregulated genes passed the p-adjusted threshold of <0.05 (p-adjusted values; *TSPY1* p = 9.72 x 10^-5^, *KDM5D* p = 0.0005, *DDX3Y* p = 0.007). GSEA enrichment from log_2_FC ranked genes produced 2 enriched and 2 downexpressed GO genesets, along with 7 enriched and 1 downregulated KEGG pathway (p-adjusted < 0.05) (**Supplementary Figure 1C**). GSEA results for GO covered upregulated Immune System process, and downregulated Response to stimulus, and multicellular organismal process. Upregulated KEGG pathways fell into major categories for Infectious Disease: Viral, Transcription, Signaling molecules and signaling, infectious disease: parasitic, and Immune disease, while the only downregulated categories was Sensory system (**Supplementary Figure 2E & 2F**). Cell deconvolution produced no significant differences between any cell signatures tested after FDR p-value correction (**Supplementary Figure 1D**).

### Effects of Spike+ Cells on the immediate immune environment

To determine the effects the spike protein has on the local immune environment, differential analysis was performed on positive and negative ROIs for Long COVID participants (**Figure 3A & 3F**). A QuPath cell classifier was created to detect SARS-CoV-2 Spike protein positive cells and statistically compare ROIs with and without positive cells (Spike+ and Spike-ROIs, respectively) for Spike protein for both colon and ileum (**Figure 3B & 3G**). In the colon, a strong statistical significance (p = 2.34x10^-10^) was found in the cellular percent frequency for positive Spike protein cells with a median of 2.09 and 0.00 for Spike protein positive and Spike protein negative ROIs, respectively. In the Ileum, a similar statistical significance was observed (p = 1.68 × 10^-8) with a median of 6.56 and 0.00 for Spike+ and Spike-ROI’s, respectively. Differential gene analysis between Spike+ and Spike-ROIs in the colon resulted in 40 downregulated (18 after BH correction) and 82 upregulated (45 after BH correction) genes with a log_2_FC > |1| and a p-value of 0.05 (**Figure 3C**). In comparison, differential analysis of the ileum showed only 10 downregulated (0 after BH correction) and 18 upregulated (0 after BH correction) transcripts (**Figure 3H**). Additionally, results were filtered based on Innate immune response (GO:0445087) and Adaptive immune response (GO:0002250) for a total of 965 and 472 transcripts, respectively, while the FC threshold was made less stringent (FC = 1.5). In the colon, 47 downregulated (1 after BH correction) and 40 upregulated (12 after BH correction) transcripts were differentially expressed in the Innate immune response while 23 downregulated (0 after BH correction) and 35 upregulated (13 after BH correction) were differentially expressed in the adaptive immune response (**Figure 3D**). Alternatively, in the ileum, 6 downregulated (0 after BH correction) and 23 upregulated (0 after BH correction) transcripts were differentially expressed for the innate immune response, while 5 downregulated (0 after BH correction) and 21 upregulated (0 after BH correction) transcripts were differentially expressed for the adaptive immune response (**Figure 3I**). Lastly, a linear regression analysis was performed for all transcripts to determine which genes have a positive or negative correlation with respect to the frequency positive cells for the Spike protein per ROI. In colon, 338 transcripts had a positive correlation (p-adjusted < 0.05) and 187 transcripts had a negative correlation (p-adjusted < 0.05) (**Figure 3E**). No transcripts were found to have a statistically significant positive or negative correlation in the ileum (**Figure 3J**).

**Figure 3.**
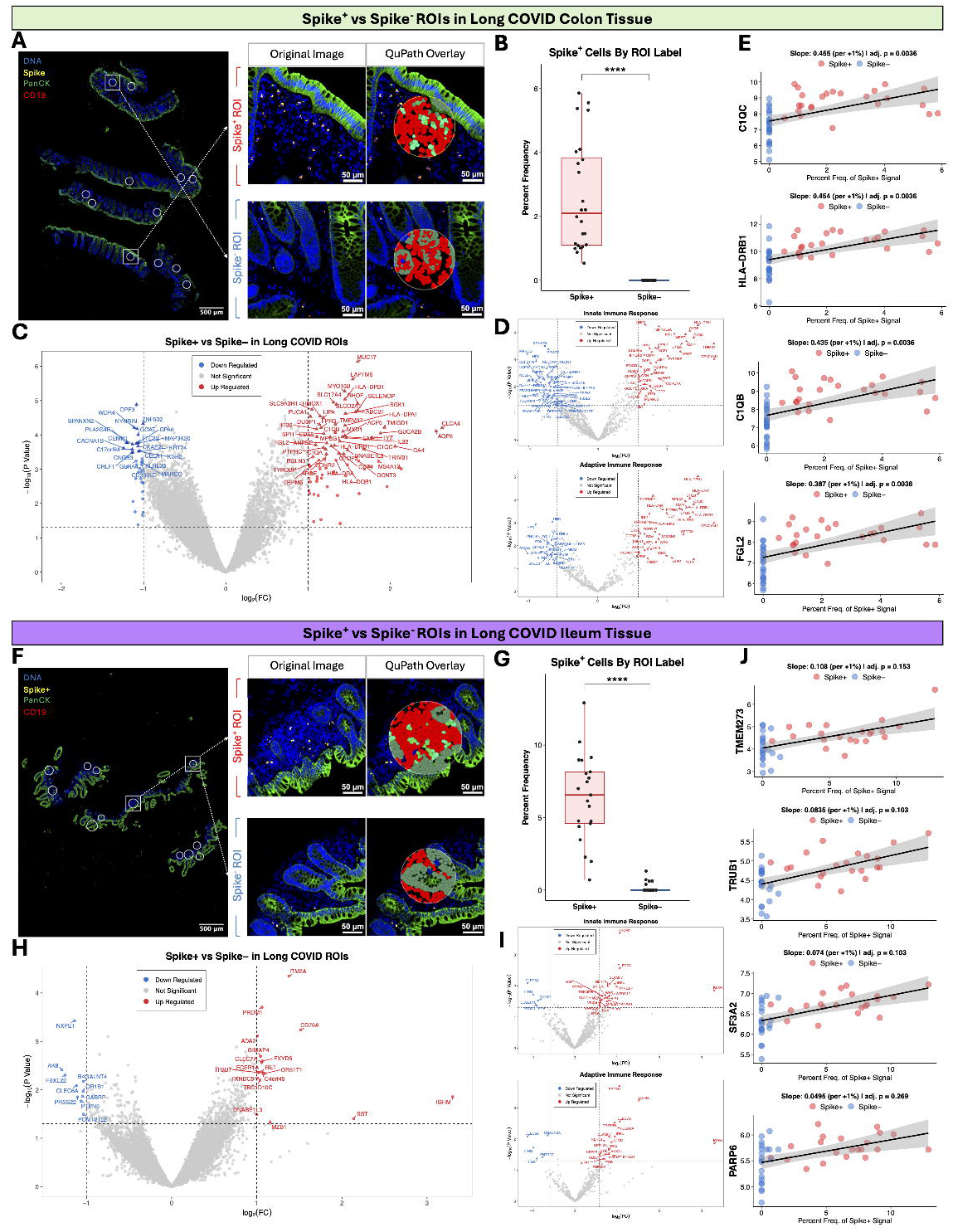
How the Presence of Spike Proteins Affects the Local Tissue Environment. (A) Immunofluorescence of a Long COVID Colon sample with GeoMx ROI’s overlayed. Zoom-In images show representative regions of Spike protein positive (Spike^+^; n = 24) and Spike protein negative (Spike^-^; n = 24) ROI’s used to create a model for differential analysis. OME.TIFF images were analyzed in QuPath, and a classification model was created to detect cells with a positive Spike signal. (B) The frequency of Spike+ cells was summarized in a box-plot representing median, min, and max. The Mann-Whitney U test found statistical significance in the frequency of detected spike antigens between Spike+ ROI’s and Spike-ROI’s in the colon (*p* = 2.34x10^-10^). (C) Differential gene analysis was performed using a limma-voom pipeline correcting for technical replicates and variability in the frequency of Spike+ cells within Spike+ ROI’s. Log_2_FC and –Log_10_(P-Value) were plotted as x-axis and y-axis, respectively. Points with a triangular shape passed the adjusted p-value threshold based on a Benjamini-Hochberg correction. 80 transcripts were upregulated (45 transcripts p-adjusted <0.05), and 40 downregulated (18 transcripts p-adjusted <0.05) (D). Sub-categories were created based on GO ontologies (GO:0002250 and GO:0045087), and the FC threshold was dropped from |2| to |1.5|; all other parameters remained the same. (E) A linear regression analysis was performed with both Spike+ and Spike-ROIs, where the x-axis represents the frequency of Spike+ Cells per ROI, and the y-axis represents the Log_2_(count) of the transcript (338 and 187 transcripts with a positive and negative slope, respectively, and p-adjusted < 0.05). Graphs shown represent the transcripts with the smallest adjusted-p value. (F) Representative immunofluorescent images of Spike+ (n = 21) and Spike- (n = 21) ROIs in Long COVID Ileum tissue. (G) The frequency of Spike+ cells was significantly different between Spike+ and Spike-ROI’s based on a Mann-Whitney U Test (p = 1.68x10^-8^) (H) Differentially expressed transcripts were plotted in a volcano with 18 upregulated and 10 downregulated (no transcripts passed the p-adjusted threshold). (I) Same description as its colon counterpart (J) No transcripts passed the p-adjusted threshold for the regression analysis. Plots shown are the top 4 transcripts with an unadjusted p-value.

Spatial deconvolution was performed using a safeTME panel to detect the transcriptional presence of immune cells in each ROI. In the colon, Spike protein positive ROIs showed a statistically significant increase in the proportion for macrophages, NC/I monocytes, plasma cells, and regulatory T cells, and a statistically significant decrease in the proportion of T CD8^+^ Naive Cells (FDR p-adjusted < 0.05) (**Figure 4A**). In comparison, the ileum showed no statistically significant difference in the proportion of each immune cell between Spike protein positive and Spike protein negative regions (**Figure 4D**). GSEA of log_2_FC-ranked genes resulted in 54 upregulated and 11 downregulated KEGG pathways and 214 upregulated and 57 downregulated GO pathways for the colon (p-adjusted < 0.05). Representative categories for KEGG and GO were selected, and the gene sets with the highest NES were plotted along with BH-corrected p-value and GeneRatio (**Figure 4B**). In KEGG, notable upregulated pathways include Antigen processing and presentation, Th1 and Th2 cell differentiation, and Th17 cell differentiation with downexpressed neuroactive ligand pathways. Notable upregulated GO pathways are multiple antigen processing and presentation gene sets, along with 3 gene sets for positive regulation of viral entry into host cell, positive regulation of viral life cycle, and non-lytic viral release. GSEA of the ileum produced 45 enriched and 9 downregulated KEGG pathways with 164 enriched and 38 downregulated GO genesets. Similar to its colon counterpart, representative categories were selected, and gene sets were plotted by NES, BH-corrected p-value, and GeneRatio (**Figure 4E**). Similar KEGG pathways appear in the ileum as in the colon, such as Th1 and Th2 cell differentiation and Th17 cell differentiation. However, some notable unique pathways are Neutrophil extracellular trap formation, COVID-19, Viral protein interaction with cytokine and cytokine receptor, and Chemokine signaling pathway. For GeneOntology, the gene sets to highlight are B cell receptor signaling pathway, respiratory burst, antigen receptor-mediated signaling pathways, and adaptive immune system. The top 50 up- and down-regulated KEGG and GO pathways for both colon and ileum were plotted to visualize pathway interactions and annotated based on the commonly occurring words and clustered (**Figure 4C & 4F**).

**Figure 4.**
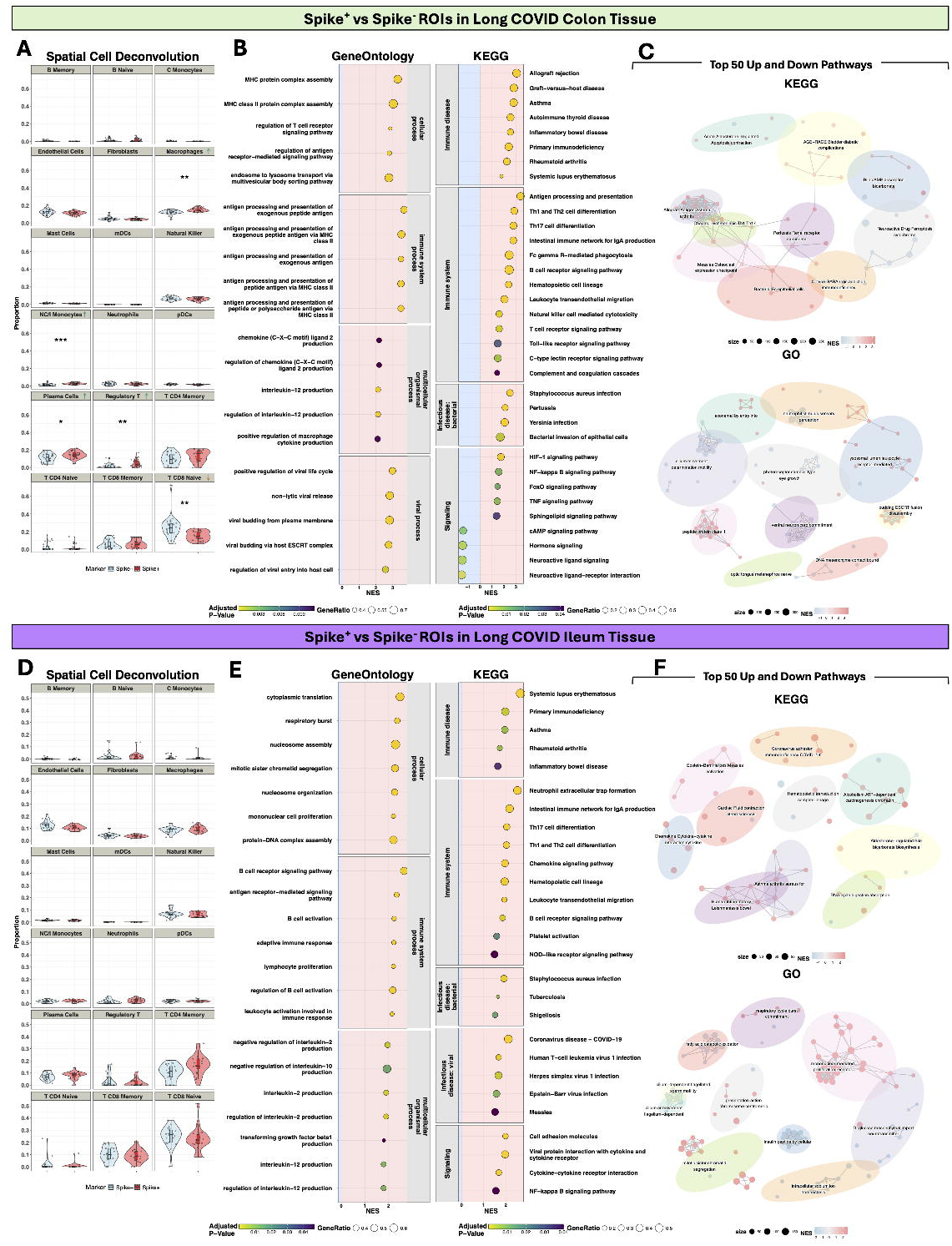
Downstream analysis of Spike Protein positive vs negative ROIs in Long COVID Tissue. (A) Spatial deconvolution was performed to compare the median cell proportion between positive Spike^+^ vs Spike^-^ using a safeTME reference expression profile. For statistically significant comparisons using the propeller framework and an FDR-corrected p-value, arrows show the direction of the Spike+ median relative to the Spike-median (FDR ≤ 0.05 (*), FDR ≤ 0.01 (**), FDR ≤ 0.001 (***)). The proportion of the combined signature of non-classical and intermediate monocytes, macrophages, regulatory T cells, and plasma cells is higher in Spike^+^, while T CD8 Naive cells are downexpressed. (B) Gene-Set Enrichment Analysis (GSEA) was performed on a ranked gene list of log^2^FC values from largest to smallest using KEGG and GO pathway and gene set references. 54 KEGG and 214 GO pathways had a NES value > 0, while 11 KEGG and 57 pathways had a NES value < 0 (p-adjusted value ≤ 0.05; GO simplified). Representative categories for KEGG and GO were selected, and pathways/ gene sets were plotted using the normalized enrichment score (NES) alongside the ratio of genes in the leading edge to total genes in the gene set (GeneRatio) and the adjusted p-value. (C) Pathways with a p-adjusted value smaller than 0.05 were selected, and the top 50 pathways for both positive (red) and negative (blue) NES values were added to the enrichment map. Clustered nodes are represented by the colored ellipses, and a group label is given based on the top 3 most frequent words in pathways. (D) Spatial deconvolution of Spike^+^ and Spike^-^ ROIs in the ileum shows no statistical significance in any comparison after p-value correction. (E) In the Ileum, 45 KEGG and 164 GO pathways had a NES value > 0 while 9 KEGG and 38 GO had a NES value < 0 (p-adjusted value < 0.05; GO simplified). (F) shares the same description as its colon counterpart.

### Multiplex Immunofluorescence of the Immune System

A custom 27-plex host immune panel was stained on serial sections of gut slides, used in spatial transcriptomics, for a small subgroup of the cohort (Long COVID n = 4 participants, healthy n = 2 participants) and processed using QuPath and R (**Figure 5A & 5B**). Representative images of the two cohorts, separated by tissue type, were grouped by complementary immune markers (**Figure 5C**). Intensity features for each marker were used to annotate similar cells based on leiden communities, and these were visualized separately between conditions for colon and ileum (**Figure 5D & 5G**). In the left colon, a total of 17 Leiden communities were annotated, with 7 communities describing the epithelium and 10 describing the connective tissue with or without immune infiltrates (**Figure 5E**). The terminal ileum had 20 leiden communities annotated with 9 belonging to the epithelium and 11 to the connective tissue. Leiden cluster identities were then exported and annotated to detected cells on QuPath and visualized onto previous field-of-views (**Figure 5F & 5I**).

**Figure 5.**
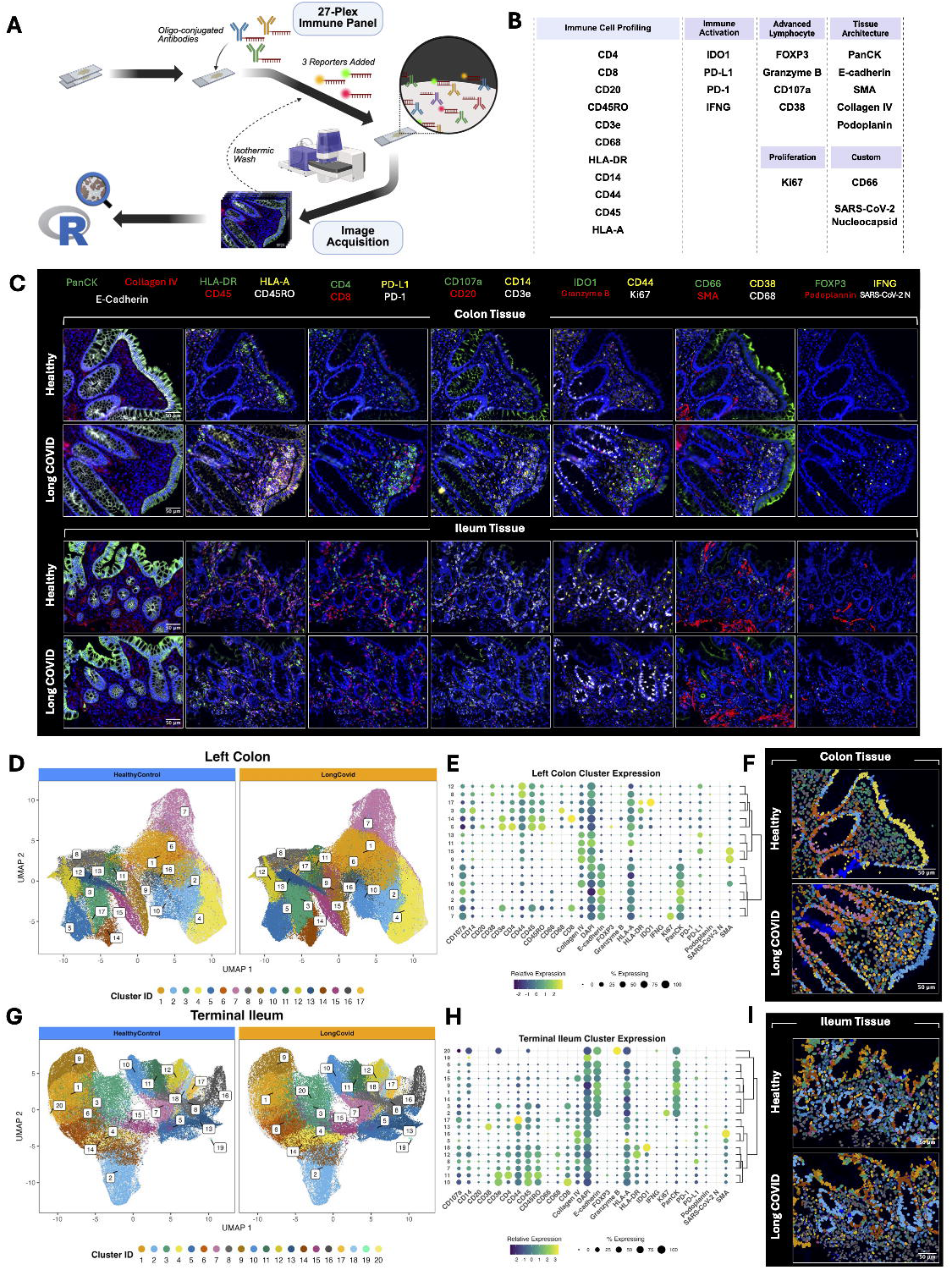
Multiplex Immunofluorescence of the Immune Environment. (A) Workflow and summary of methods. (B) List of antibody targets for the study (Total targets = 27). (C) Four representative images processed in QuPath and separated by markers comparing Long COVID samples to Healthy in both the Ileum and Colon. (D) After cell detection and feature extraction, single-cell data from the colon were processed in R using a pipeline to detect Leiden clusters from z-score normalized cell expression data and plotted on a UMAP separated by condition. A total of 17 clusters were detected for the colon tissue. (E) The dot plot compares marker expression via two metrics throughout all 17 clusters and then is hierarchically organized to highlight marker expression patterns. The two metrics measured are “% expressing” or percent of positive cells out of all the cells in the cluster, and “Relative expression” or the average cell MFI transformed to a z-score and scaled to each marker. The first major split in the hierarchy occurs between clusters belonging to the epithelium versus stromal compartments, followed by immune versus non-immune groups. (F) Cluster identity was annotated to each cell and exported back to QuPath to visualize detected clusters within the tissue. (G) The same processing pipeline was performed in the ileum, independent of the colon data. In the ileum, 20 clusters were detected. (H) In the ileum, the first hierarchy distinction occurs between epithelium and stromal cells, but the epithelium is denoted by an excess cluster characterized by cells expressing Granzyme B in the epithelium. (I) Shares the same description as its colon counterpart.

## Discussion

Our findings establish that SARS-CoV-2 Spike RNA and protein persist in the gastrointestinal (GI) tissue of LC patients long after acute infection resolves. Crucially, this persistence is not immunologically inert; it is associated with a localized immune microenvironment with distinct functional transcriptional programs that are not observed in healthy tissue. These results align with the hypothesis that retained viral RNA and/or antigens act as chronic stimuli, sustaining chronic immune activation and preventing inflammatory resolution.

### Transcriptional profile of Spike+ ROIs in LC tissue vs. controls

We first compared transcriptional profiles in Spike+ regions of LC tissue versus Spike+ regions from controls. SARS-CoV-2 Spike transcript (*nCoV2019-S*) and Spike protein were identified in the colon and ileum tissue of LC patients and controls using RNAscope and antibody staining. Surprisingly, a participant with no recorded infection was positive for *nCoV2019-S* and Spike protein. This observation could be explained by a prior asymptomatic or otherwise unrecognized infection that was not clinically recorded. Several studies estimate a substantial proportion of SARS-COV-2 infections are asymptomatic, with many reports suggesting approximately 35 - 40% of confirmed infections occur without symptoms.^35,36^ As a result, individuals may have been infected without undergoing testing or receiving a clinical diagnosis, since to many the absence of symptoms provided little reason to seek testing.^37^ This outlines the need for pre-pandemic controls to ensure that participants tested were not previously infected with SARS-CoV-2. Nonetheless, comparison of the immune microenvironment in Spike+ tissue regions within colon tissue from cases vs. controls showed significant differences in gene expression changes near the Spike protein. These differences are not found when comparing the entire tissue or bulk analysis of RNA, it requires specialized region of interest comparison to detect these new gene dysregulations.

These transcriptional changes extend beyond immune activation to disrupt tissue homeostasis. For instance, we observed a simultaneous downregulation of key homeostatic chemokines, such as *CXCL13*, *CCL19,* and *CCL21*. This paradox—intense local inflammation alongside suppressed trafficking signals—points to disrupted immune surveillance or a localized “exhaustion” phenotype. Interestingly, these three genes, along with *DMBT1, MS4A1* (CD20), and *C3*, drive the leading edge of the downregulated GeneOntology gene set for humoral response. An uncoordinated or downexpressed humoral response has been observed in other LC studies that looked at blood or serum.^38,39^ A late 2025 study, from Beth Israel Deaconess Medical Center, transcriptionally compared peripheral blood of LC versus convalescent controls and found an upregulation of multiple cytokines, including *IL-6*, along with a downregulation of T and B cell signatures in their 2023-2024 cohort of samples.^40^ In our study, *IL-6* expression RNA transcript, and not the soluble cytokine, trends were downregulated (did not meet the Log_2_FC threshold), but the dysregulated B cell phenotype was present. This highlights the heterogeneity of the disease and could imply the disease is characterized differently between blood/ serum and tissue. Future studies should directly compare immunological responses in matched blood and tissue samples from the same LC participants and controls.

Upregulation of the water-channel gene AQP8 and absorptive genes (*SLC26A3*, *SLC26A2*) in affected regions represents a compensatory response to epithelial stress. Because aquaporin dysregulation is frequently linked to gastrointestinal distress, this could provide a molecular basis for the chronic GI symptoms (e.g., diarrhea and abdominal pain) reported by Long COVID patients. However, studies report that *AQP8* expression varies between disease and this gene has not previously been associated with Long COVID.^41,42^ A study from 2023 on inflammatory bowel disease (IBD) found that *AQP8* and *SLC26A2* (along with 8 other genes) potentially contribute to tissue repair and *AQP8* was associated with improved recovery of bowel function.^43^ Multiple human and mouse studies have found that *SLC26A3* expression contributes to a healthy intestinal epithelial layer and its loss contributes to IBD and ulcerative colitis.^44–46^ This could be an indication that the epithelial barrier of the LC colon is in process of repair and possibly explain upregulated myofibroblast differentiation, microvilli organization, and cell-to-cell adhesion signaling genesets.

Overall, the findings indicate that although Spike protein can be identified in tissue from LC cases and controls, the ongoing symptomatic LC state is associated with differential local immune responses compared to controls. This suggests that LC patients have some form of pre-existing vulnerability that sets them up to have a different immunological response to persistent Spike protein. This could be a genetic vulnerability, or perhaps dysbiosis of the gut microbiome environment and/or low-level subclinical inflammation could predispose to altered Spike protein. Further research is needed that tests how the gut microbiome or other medical comorbidities affect the host’s response to the Spike RNA or protein.

### Transcriptional profiles of Spike+ vs. Spike-ROIs in LC

Our second analysis compares Spike+ vs. Spike-ROIs in LC to answer the question of what type of cellular environment the Spike protein may be found. Cellular deconvolution of Spike+ ROIs reveals enrichment of myeloid-derived cells (intermediate/non-classical monocytes and macrophages), plasma cells, and T-regulatory cell signatures. Our linear regression analysis compliments these results by highlighting a positive correlation between increasing Spike protein count and gene counts involved in antigen processing, the classical complement pathway, and inflammation. Together, upregulation of antigen-processing programs and focal enrichment of macrophages, intermediate/non-classical monocytes, and plasma cells within Spike+ regions demonstrate active, sustained immune engagement with persistent antigen.

A 2021 autopsy study found that, during acute infection, cells that were the most enriched for SARS-CoV-2 RNA in the lung were *CD14^high^CD16^high^*monocytes and *LDB2^high^OSMR^high^YAP1^high^* macrophages.^16^ This finding complements our statistically significant differences in gene signatures for intermediate/ non-classical monocytes and macrophages. RNAscope images of the Long COVID colon produced in Peluso et. al. show *Spike* gene signal found within cells of the lamina propria and minimal signal reported in the epithelium similar to the images we present.^24^ Their IHC staining of subsequent sections where *Spike* signal was found showed positive staining for CD68+ and, rarely, CD3+ cells. Our results did not indicate a differential expression of any gene part of the *CD3* family, albeit deconvolution did show increased T-reg signatures. However, our study shows *CD68* was found to be upregulated in Spike+ ROIs adding to the mounting evidence that Spike RNA and protein can localize near monocytes/ macrophages.

The data presented here suggests SARS-CoV-2 Spike RNA or protein can be taken up by myeloid cells that aren’t involved via canonical ACE2-mediated viral entry in Long COVID. This is supported by a study from 2024 which showed that a SARS-CoV-2 Spike-pseudotyped lentivirus was able to infect human primary interstitial macrophages in the lung that do not express ACE2, but infection was prevented using DC-SIGN/CD209 blocking antibodies.^47^ In addition, work by Livanos et. al. provides context of how the acute viral infection of the small intestine occurs through the immunofluorescent detection of SARS-CoV-2 nucleocapsid (N) protein in ACE2 expressing intestinal epithelial cells.^17^ A plausible mechanism is that the virus first infects ACE2-expressing parenchymal cells, after which myeloid cells uptake the viral RNA or protein and contribute to the viral persistence. Future studies in LC should characterize and test this mechanism directly. However, the study we present reports that dysregulated or exhausted antigen processing and presentation could be a strong lead in explaining why viral spike protein can persist in some tissues.

The concept that pathogens can reside in situ within tissue, persisting not as primary aggressors but as opportunists enabled by host immune insufficiency, is well established across diverse infectious diseases and provides critical context for interpreting our findings. Cytomegalovirus (CMV), a ubiquitous herpesvirus carried in latent form by the majority of adults, is a paradigmatic example: it persists silently in tissue-resident myeloid cells and endothelial cells but reactivates and drives overt disease specifically when host immune surveillance is impaired through immunosuppression, transplantation, or immune exhaustion.^48^ Immune control of CMV reactivation has been shown to depend critically on functional T cell responses; its disruption permits the virus to escape from latency and establish pathogenic tissue infection.^49^ Similarly, Epstein-Barr virus (EBV) establishes lifelong latency in mucosal tissues and B lymphocytes, forming ectopic lymphoid structures in various organs.^50^ In the context of Long COVID and conditions such as myalgic encephalomyelitis/chronic fatigue syndrome, EBV reactivation has been proposed to emerge from a state of acquired immune deficiency, wherein EBV-specific CD4+ T cell surveillance fails, permitting latent viral programs to re-engage, drive chronic inflammation, and compound immune exhaustion.^51^ Critically, both EBV^52–54^ and CMV^55^ reactivation have been documented in acute COVID-19 and post-acute sequelae, suggesting that SARS-CoV-2 itself generates a permissive immunological environment that allows co-resident latent pathogens to thrive in tissue. Beyond viral pathogens, bacteria that reside in gut mucosal tissue also follow this host-deficiency-driven paradigm. Adherent-invasive *Escherichia coli* (AIEC) strains, abnormally enriched in the intestinal mucosa of Crohn’s disease patients, invade intestinal epithelial cells and exploit defective autophagy and impaired innate immunity to survive within macrophages.^56,57^ The ability of AIEC to persist intracellularly is not a function of bacterial virulence alone, but depends on host cellular deficiencies — particularly in autophagy pathways — that compromise bacterial clearance. Similarly, *Helicobacter pylori* establish a remarkably durable in situ colonization of the gastric mucosa by manipulating the host immune landscape, suppressing effector T cell responses, and creating an immunosuppressive tissue microenvironment that mirrors the exhaustion phenotypes we observe in LC gut tissue. Single-cell transcriptomic analysis of H. pylori-infected gastric mucosa has revealed HLA-DR+ and CTLA4-expressing T cell populations consistent with immune suppression and antigen-driven exhaustion, allowing the bacterium to persist for decades in tissue.^58^ Collectively, these examples underscore a unifying principle: pathogens residing *in situ* in tissue are not inherently unstoppable, but rather, it is the failure of host immune surveillance that allows them to persist, replicate, and drive chronic pathology. Our findings of immune exhaustion signatures, impaired lymphocyte trafficking, and dysfunctional myeloid antigen presentation in LC gut tissue are consistent with this framework: the host immune deficiencies we document are not merely a consequence of viral persistence but are likely a precondition — or an actively maintained state — that prevents resolution of SARS-CoV-2 antigen from tissue. Targeting these host-intrinsic vulnerabilities, rather than the pathogen alone, may therefore represent a more tractable therapeutic strategy.

### The Spike Protein as a Catalyst for Tumorigenesis

We identified enriched gene sets associated with tumorigenesis within Spike+ niches. While these signatures do not indicate active malignancy, they suggest that the chronic inflammatory environment induced by persistent Spike protein promotes cellular stress and proliferative signaling pathways mirroring other known pre-cancerous states. This observation gains importance when viewed through the lens of current epidemiological shifts. According to recent American Cancer Society reports^59,60^, colorectal cancer (CRC) has unexpectedly emerged as the leading cause of cancer mortality in adults under age 50. This 2% annual increase in CRC mentioned in recent reports provides a critical clinical backdrop for your finding that Spike+ regions in the colon exhibit transcripts specifically associated to tumorigenesis and progression, such as *GUCA2A*, *TSPAN1,* and *S100P*.^61–63^ Although these genes aren’t known to contribute directly to CRC, their dysregulated expression could support a tissue environment that is more permissive for tumorigenesis. While lifestyle factors are contributory, they do not fully account for the sharp 2% annual increase in early-onset CRC observed over the last decade — a trend that appears to have accelerated in the post-pandemic era.

However, we also noted that known tumor suppressor genes are also enriched in LC such as *SLC26A3* and *CLCA4*.^64,65^ The concurrent presence of both tumor-promoting and tumor-suppressive transcriptional programs suggests that Spike+ niches do not represent a malignant state, but rather a dynamic microenvironment characterized by competing regulatory signals. This may reflect a compensatory epithelial response to chronic injury and inflammation in which stress-induced pathways are activated alongside mechanisms that attempt to preserve or return to tissue homeostasis. Whether this transcriptional landscape confers increased long-term susceptibility to neoplastic formations, or instead represents a reversible inflammatory adaptation, remains unknown. Longitudinal studies integrating spatial transcriptomics with clinical follow-up will be required to determine whether these localized molecular signatures translate into measurable changes in CRC risk.

### Tissue-Specific Vulnerability and Limitations

The detection of SARS-CoV-2 Spike transcript (*nCoV2019-S*) and the Spike protein were identified in the colon and ileum tissue of both LC patients and controls. The small sample size of our control cohort constrains our confidence to say all healthy individuals carry persistent Spike RNA and/or protein (n = 8 Long COVID, n = 5 Controls). Additionally, previously published colon images with negative *Spike* RNA, by Peluso et. al, come from pre-pandemic control samples which our study did not look at.^24^ Future studies should examine tissue collected from at larger control cohorts from both pre- and post-pandemic samples to confirm whether viral RNA and protein can be found in a healthy post-pandemic cohort. A second limitation is that the RNA probes target SARS-CoV-2 Spike and orf1ab from the original Wuhan strain. Other SARS-CoV-2 RNA and protein targets (including strains) or subgenomic RNA in the lower GI tract should be targeted and detected with alternative methods such as ddPCR or metagenomic sequencing. The small sample size limits generalizability to the broader LC population. Given that the LC cohort was predominantly female and the control cohort predominantly male, sex-associated transcriptional differences may contribute to the observed differential expression and should be considered when interpreting these findings. Additionally, many of our LC participants have a history of other medical conditions and actively take medications that could have an impact on the transcriptome of the small and large intestine. Therefore, future LC omics studies must enroll larger, well-matched cohorts to validate these findings and identify factors predisposing patients to altered immunological responses to gut-resident Spike RNA or protein.

Observed transcriptional changes in the LC left colon could be explained by pre-existing vulnerabilities that determine how the local immune environment responds to the presence of Spike RNA or protein that differs from the terminal ileum. Studies comparing the immune cell composition between the left colon and terminal ileum outline regional differences that shape and contribute to different immune cell populations and activation states, which may explain why these tissues respond differently to inflammatory stimuli even within the same patient.^66–68^ These immune differences may contribute to region-specific manifestations of inflammatory bowel diseases (IBD) by creating niche conditions that favor the initiation or amplification of inflammatory responses. The colon also harbors a larger bacterial load than the ileum, and healthy individuals harbor distinct microbiomes between the two regions fueling the idea that different immune system functions are needed to maintain gut homeostasis.^69^

Comparison of the immune microenvironment in Spike+ tissue regions of the colon highlights distinct and significant gene expression changes near the Spike protein. Pathway-level analysis further demonstrated enrichment of immune-related signaling pathways in the left colon of LC patients compared to controls that are not detected in the terminal ileum. Transcriptional profiles of Spike+ vs. Spike-ROIs in LC suggest that the colon may be a primary “hotspot” for post-viral immunopathology, a finding that should inform future biopsy-based diagnostics and therapeutic targeting. In summary, our data discusses a model in which persistent SARS-CoV-2 Spike protein in the gut establishes a localized, pro-inflammatory niche characterized by myeloid dysfunction and altered epithelial homeostasis. These findings provide a direct biological mechanism for how viral-induced inflammation drives the “accelerated aging” and rising cancer rates observed in the post-COVID era. Targeting these viral reservoirs represents a promising therapeutic strategy to prevent the long-term systemic and neoplastic sequelae of COVID-19.

## Acknowledgments

We are grateful to the staff of the La Jolla Institute of Immunology Histology Core and the JCVI Genomic Core for their continuous support in tissue processing, imaging, and sequencing that made this work possible. We also thank Amy Proal, Michael VanElzakker, Sunny Jang, and Eric Weine for their contributions to this study.

## Author Contributions

SM, DP, and MF conceptualized the study. SA was responsible for data curation and investigation. SA and PP did the formal data analysis. SA did the laboratory work. MF oversaw the project and acquired funding. SA and MF wrote the initial draft of the manuscript. All authors reviewed and edited the manuscript. The corresponding author had full access to all the data in the study and had final responsibility for the decision to submit for publication.

## Code Availability

R code used for computational analysis will be made available on GitHub (https://github.com/drfreire).

