## Supplementary Figure for "Persistent SARS-CoV-2 Spike is Associated with Localized Immune Dysregulation in Long COVID Gut Biopsies"

### Supplementary Figures

**Supplementary Table 1.**

| Variable | Control N = 5[1] | Long COVID N = 8[1] | p-value[2] |
| --- | --- | --- | --- |
| **Tissue Collected** | | | |
| Colon & Ileum | 5 / 5 (100%) | 7 / 8 (88%) | |
| Colon Only | 0 / 5 (0%) | 1 / 8 (13%) | |
| **Age** | 48 (18) | 45 (12) | 0.94 |
| **Sex** | | | 0.10 |
| Female | 1 / 5 (20%) | 6 / 8 (75%) | |
| Male | 4 / 5 (80%) | 2 / 8 (25%) | |
| **BMI** | 28.2 (4.1) | 24.4 (5.0) | 0.12 |

[1] n / N (%); Mean (SD)

[2] Wilcoxon rank sum test; Fisher’s exact test

**Supplementary Table 1.** Demographics of Patients by Condition

### Supplementary Figure 1.

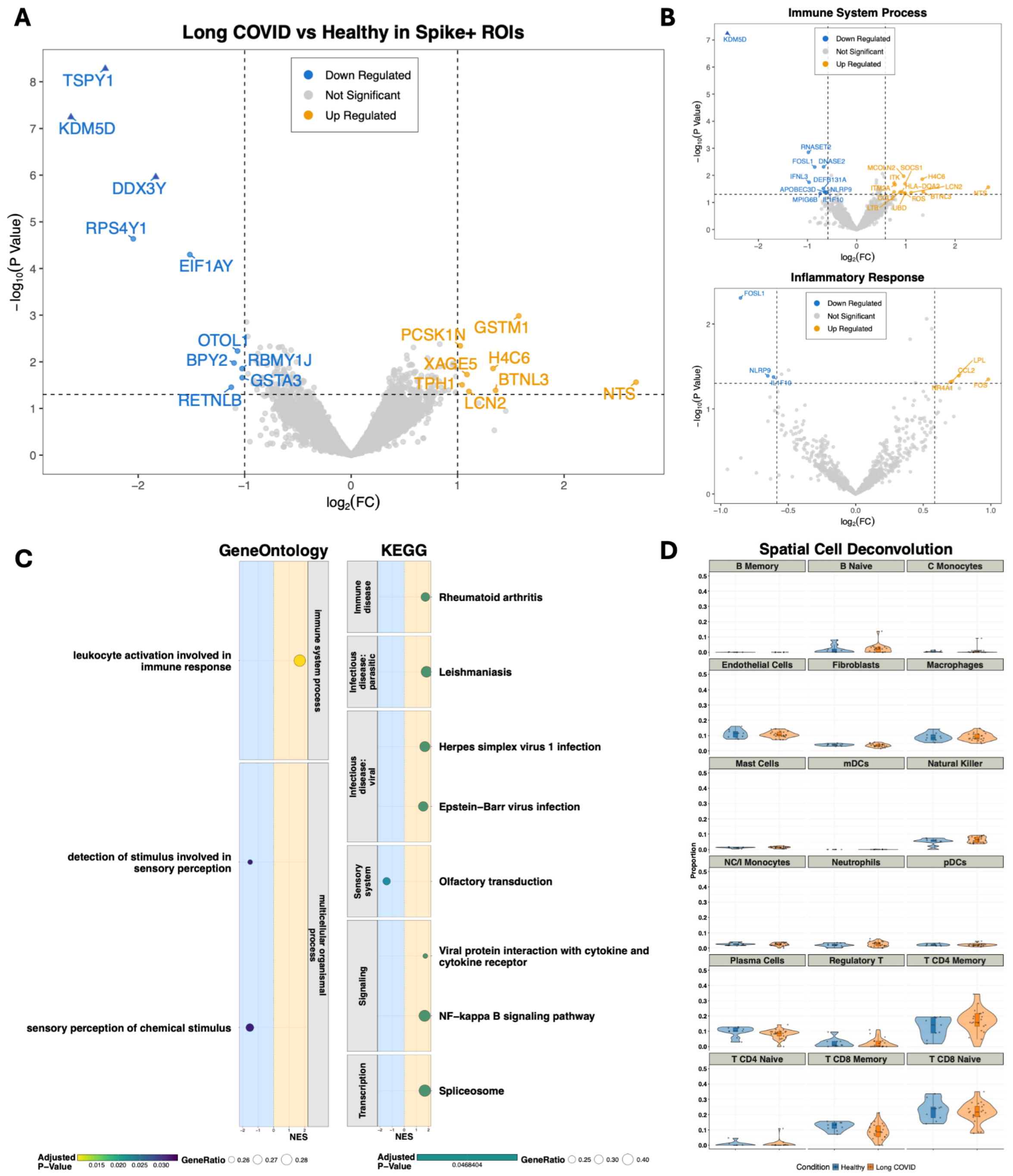

**Supplementary Figure 1. Differential Expression of Long COVID and Healthy Spike+ ROIs in the Ileum by Spatial Transcriptomics.** (A) A total of 33 ROIs were analyzed between 8 Long COVID (24 ROIs) and 3 healthy (9 ROIs) participants in the region positive for SARS-CoV-2 Spike antigen. Differential analysis was performed using the limma-voom pipeline, correcting for replicate samples and variation in the frequency of detected SARS-CoV-2 Spike by ROI. 18,581 transcript targets were plotted. The main volcano plot highlights 8 and 10 genes that are upregulated and downregulated, respectively, at an unadjusted *p*-value threshold of 0.05 and an absolute $\log_2$FC > 1. Points with a triangular shape passed the adjusted p-value threshold based on a Benjamini-Hochberg correction. (B) Genes involved in *GeneOntology* (GO) (GO:0002376 & GO:0006954) parent category were filtered and plotted into each respective volcano plot with an unadjusted *p*-value threshold of 0.05 and an absolute FC > 1.5. (C) Gene-Set Enrichment Analysis was performed on a ranked list of $\log_2$FC values, resulting in 4 *GeneOntology* gene sets (2 enriched and 2 downexpressed) and 8 *KEGG* pathways (7 enriched and 1 downexpressed) passing the adjusted p-value threshold of 0.05. Representative categories were selected, and pathways/ gene sets were plotted using the normalized enrichment score (NES) alongside the ratio of genes in the leading edge to total genes in the gene set (GeneRatio) and the adjusted p-value. (D) Spatial deconvolution was performed using a safeTME reference expression profile. Statistical significance was assessed using the propeller framework, a moderated two-tailed Mann-Whitney U test with BH FDR correction (*FDR* ≤ 0.05 (*), *FDR* ≤ 0.01 (**), *FDR* ≤ 0.001 (***)). No statistical significance was found in any cell type compared.

### Supplementary Figure 2.

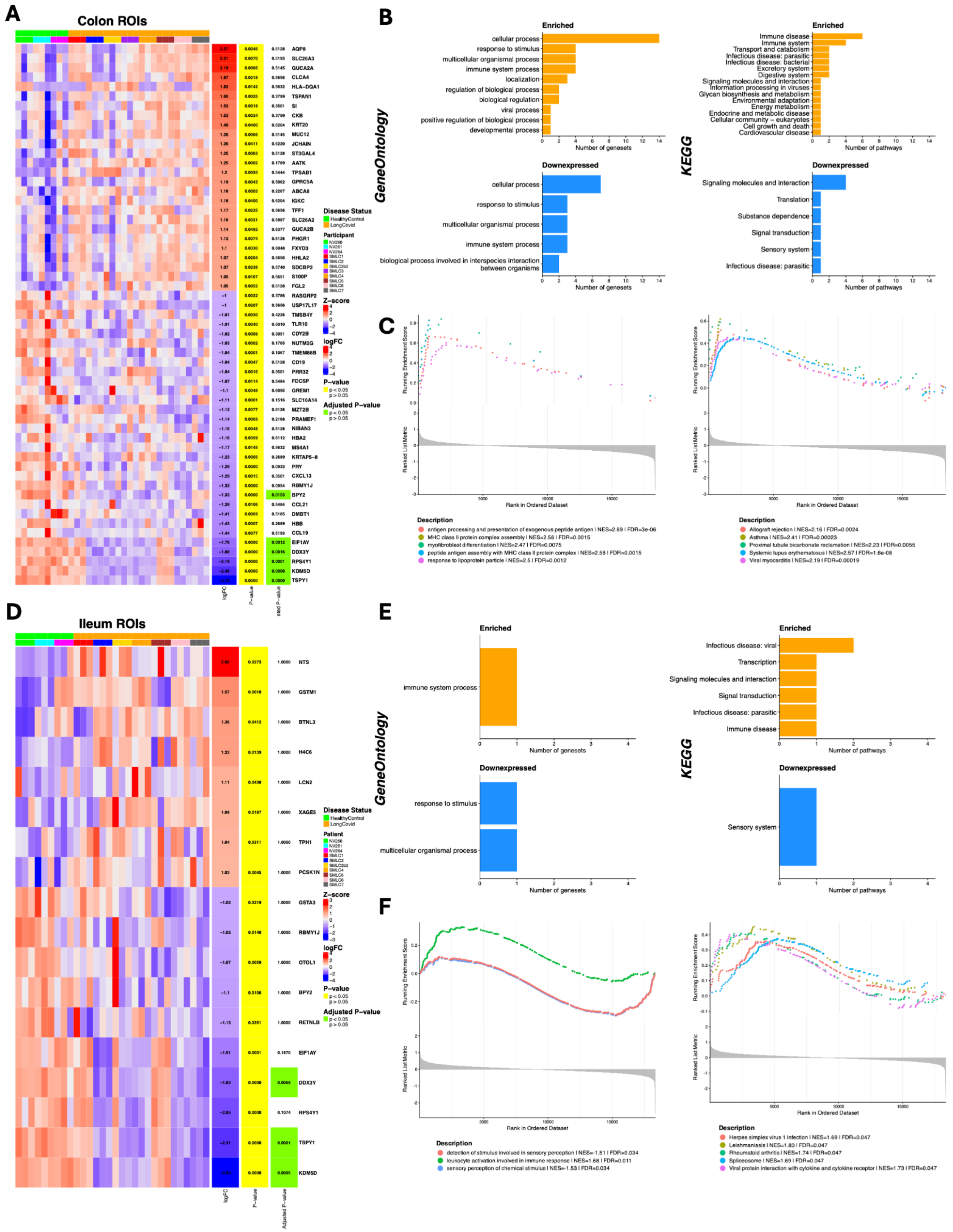

**Supplementary Figure 2. GSEA enrichment hierarchy count and top leading-edge.** (A) (D) heatmap of differentially expressed genes (by p-value < 0.05) showing row-wise z-scored log-normalized expression scores organized by condition then participant. (B) (E) Count of genesets/ pathways in GeneOntology (BP) and KEGG by major category produced from GSEA (BH adjusted p-value < 0.05). (C) (F) The top 5 pathways by absolute NES were selected and plotted in a leading-edge enrichment plot with rank list metrics.

**Supplementary Table 2.**

| Patient ID | Condition | Comorbidities | Medications |
|---|---|---|---|
| SMLC001 | Long COVID | hx of Long COVID, allergic rhinitis, hyperlipidemia, vitamin D deficiency, colon polyps | aspirin, lactobacillus 3/FOS/pantethine, quercetin, alpha-lipoic acid, American ginseng root, bromelains, cholecalciferol, coenzyme Q, famotidine, fexofenadine, levocarnitine, montelukast, omega 3-dha-epa-fish oil, turmeric, vitamin B complex capsule |
| SMLC002 | Long COVID | hx of Long COVID, appendicitis, GERD, Hashimoto's thyroiditis, hemorrhoids, colon polyps | rimegepant, nebivolol, candesartan, ondansetron, spironolactone, levothyroxine |
| SMLC003 | Long COVID | hx of Long COVID, dysautonomia | none |
| SMLC004 | Long COVID | hx of Long COVID, proctitis/periappendical inflammation | none |
| SMLC002B2 | Long COVID | hx of Long COVID, appendicitis, GERD, Hashimoto's thyroiditis, hemorrhoids, colon polyps, mildly active nonspecific colitis found on 7/27/2023 | rimegepant, nebivolol, candesartan, ondansetron, spironolactone, levothyroxine |
| SMLC005 | Long COVID | hx of Long COVID, hemorrhoids | acetylcysteine, cyanocobalamin, guanfacine, pyridoxine, pyridostigmine, valganciclovir, finasteride |
| SMLC006 | Long COVID | hx of Long COVID, hemorrhoids, anxiety, depression | none |
| SMLC007 | Long COVID | hx of Long COVID, PCOS, severe obesity, menorrhagia, oligomenorrhea, PTSD, ADHD, dysautonomia | tizanidine, gabapentin, propranolol, cholecalciferol, alprazolam |
| NV260 | Control | hx of anxiety, low blood pressure, thyroid disease | none |
| NV261 | Control | hx of heart failure due to valvular disease, hyperlipidemia, hypertriglyceridemia | none |
| NV264 | Control | hx of dyslipidemia, vitamin D deficiency, peptic duodenitis | none |
| NV362 | Control | hx of colon polyps | famotidine, Valsartan-Hydrochlorothiazide |
| NV363 | Control | hx of colon polyps, breast cancer, hypertension, osteoarthritis | memantine, losartan |

**Supplementary Table 2.** Demographics by Patient annotated with Comorbidities and Medications.

**Supplementary Table 3.**

| Patient ID | Condition | Date of Biopsy | Date of Last Infection | Vaccination History |
|---|---|---|---|---|
| SMLC001 | Long COVID | 11/11/22 | 4/16/22 | COVID-19 MODERNA 12Y+, 0.5 mL - 10/23/2021, 05/28/2022<br>COVID-19, Ad26, 0.5mL (Janssen) (Johnson and Johnson Covid-19 Vaccine EUA) - 4/2/2021 |
| SMLC002 | Long COVID | 11/15/22 | 2/8/21 | 2vCOV-mRNA, 30mcg/0.3mL, 12y+ (Pfizer) - 10/14/2022<br>1vCOV-mRNA, 30mcg/0.3mL, 12y+ (Pfizer) - 12/22/2021, 4/21/2021, 3/31/2021 |
| SMLC003 | Long COVID | 3/28/23 | ~07/22 (reported) | No vaccine record. |
| SMLC004 | Long COVID | 5/23/23 | ~02/22 (reported) | 2vCOV-mRNA, 30mcg/0.3mL, 12y+ (Pfizer) - 9/23/2022<br>COVID-19 Vaccine - PFIZER - 03/19/2021, 02/26/2021 |
| SMLC002B2 | Long COVID | 7/27/23 | 2/8/21 | 2vCOV-mRNA, 30mcg/0.3mL, 12y+ (Pfizer) - 10/14/2022<br>1vCOV-mRNA, 30mcg/0.3mL, 12y+ (Pfizer) - 12/22/2021, 4/21/2021, 3/31/2021 |
| SMLC005 | Long COVID | 10/23/23 | 1/7/22 | COVID-19, mRNA, 2023-24, 2024-25, 30mcg/0.3mL, 12y+ (COMIRNATY) - 9/29/2023<br>2vCOV-mRNA, 30mcg/0.3mL, 12y+ (Pfizer) - 9/9/2022<br>1vCOV-mRNA, 30mcg/0.3mL, 12y+ (Pfizer) - 2/25/2021, 03/19/2021, 09/29/2021 |
| SMLC006 | Long COVID | 10/23/23 | ~07/22 (reported) | Covid-19 Vaccine, Moderna - 11/24/21<br>Covid-19 Vaccine, Janssen - 03/01/21 |
| SMLC007 | Long COVID | 12/7/23 | 6/9/22 | 1vCOV-aPS, NVX-CoV2373, 0.5mL (Novavax) - 4/25/2023<br>1vCOV-mRNA, 30mcg/0.3mL, 12y+ (Pfizer) - 12/16/21 |
| NV260 | Control | 6/20/23 | 5/18/22 | 1vCOV-mRNA, 30mcg/0.3mL, 12y+ (Pfizer) - 02/17/2021, 01/27/2021, 12/18/2021 |
| NV261 | Control | 6/20/23 | NA | 1vCOV-mRNA, 100mcg/0.5mL (Moderna) - 2/6/2021, 1/9/2021 |
| NV264 | Control | 7/25/23 | NA | 1vCOV-mRNA, 30mcg/0.3mL, 12y+ (Pfizer) - 12/8/2021 |
| NV362 | Control | 6/26/25 | NA | No vaccine record. |
| NV363 | Control | 6/26/25 | ~03/20 (reported) | 1vCOV-mRNA, 30mcg/0.3mL, 12y+ (Pfizer) - 9/27/2021, 2/24/2021, 2/3/2021 |

**Supplementary Table 3.** Demographics by Patient annotated with Date of Last COVID-19 Infection and Vaccination History.

### Supplementary Figure 3.

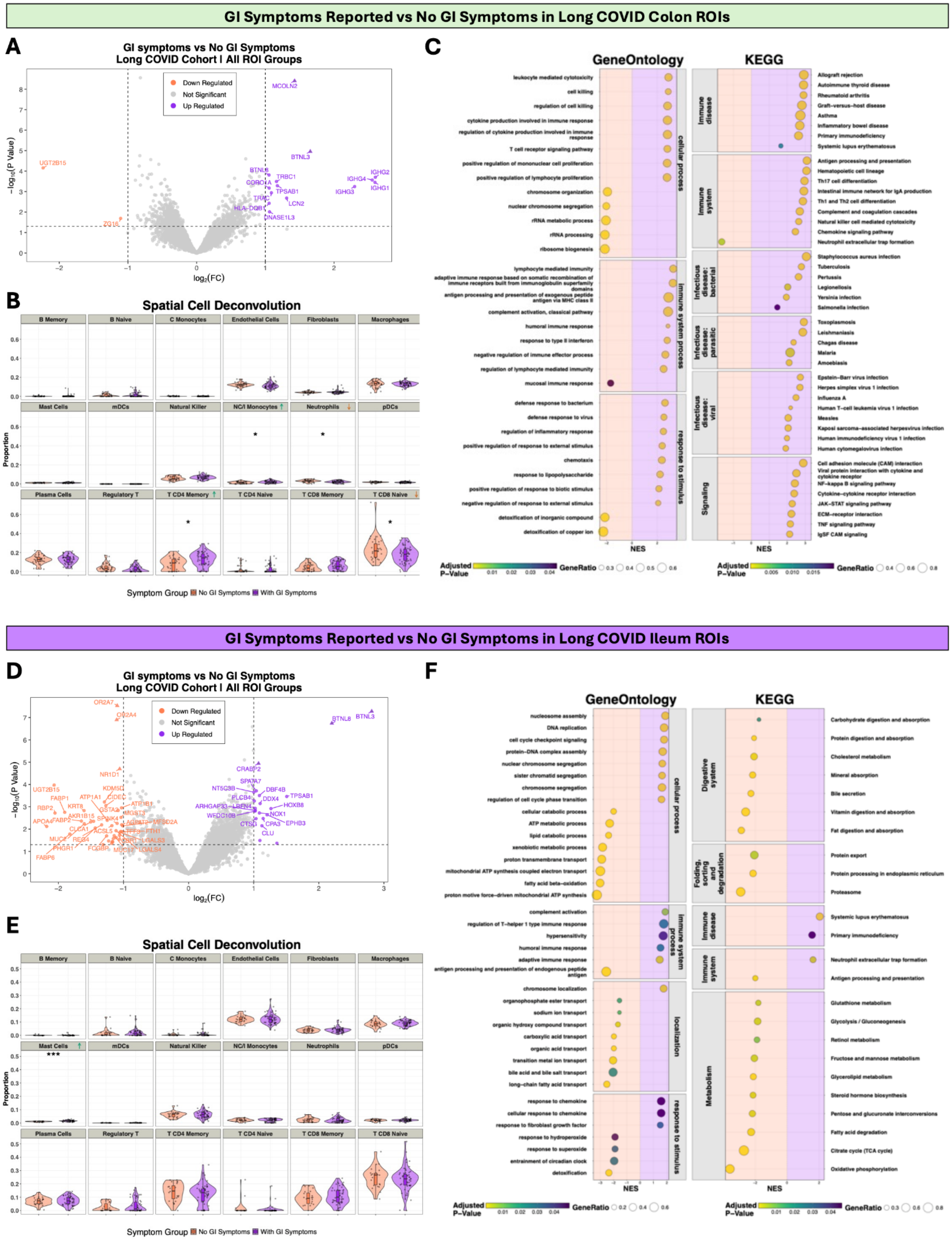

**Supplementary Figure 3. Differential Analysis of the Colon and Ileum between Long COVID Patients with and Without GI Symptoms.** (A) Differential expression of Colon ROIs for Long COVID participants with GI symptoms (5 patients) and without GI symptoms (3 patients) produced 14 upregulated and 2 downregulated statistically significant genes. (B) Spatial cell deconvolution was performed using safeTME as the reference dataset; no statistically significant (BH adjusted p-value < 0.05) comparisons were found. (C) GSEA was performed on a ranked list of $log_2FC$ values, resulting in 341 *GeneOntology* genesets (250 enriched and 91 downexpressed) and 119 *KEGG* pathways (86 enriched and 33 downexpressed) passing the BH-adjusted p-value threshold of 0.05. Representative categories were selected, and pathways/ gene sets were plotted using the normalized enrichment score (NES) alongside ratio of genes in the leading edge to total genes in the gene set (GeneRatio) and the adjusted p-value. (D) Differential expression of Ileum ROIs for Long COVID participants with GI symptoms (4 patients) and without GI symptoms (2 patients) produced 20 upregulated and 40 downregulated statistically significant genes. (E) Spatial cell deconvolution was performed using safeTME as the reference dataset; no statistically significant (BH adjusted p-value < 0.05) comparisons were found. (F) GSEA was performed on a ranked list of $log_2FC$ values, resulting in 171 *GeneOntology* genesets (55 enriched and 116 downexpressed) and 70 *KEGG* pathways (10 enriched and 60 downexpressed) passing the adjusted p-value threshold of 0.05.
